# FIRST EVIDENCE OF PRESENCE OF *RALSTONIA SOLANACEARUM* IN *THRIPS HAWAIIENSIS* (THYSANOPTERA: THRIPIDAE) IN INDANG, CAVITE

**DOI:** 10.64898/2026.07.16.738829

**Authors:** Alliah Czarielle N. Mañugo, Jay-Vee S. Mendoza, Jacinth M. Jungco, Rizalina L. Tiongco, Bonn Andrei C. Revilleza, Rachele L. De Torres, Oliver D. Balanban, Fe M. Dela Cueva

## Abstract

Bugtok disease remains a major bacterial constraint of cooking banana production in the Philippines and is characterized by vascular discoloration, fruit browning, and progressive decline associated with members of the *Ralstonia solanacearum* species complex that infect banana inflorescences and fruits. This study investigated whether insects visiting *Saba* banana flowers in Indang, Cavite harbor *R. solanacearum*, with emphasis on the possible role of *Thrips hawaiiensis* as a possible candidate vector under field conditions. Destructive sampling was conducted three times in a Saba-monoculturing farm with high reported bugtok incidence, targeting flowers present at the time of collection and prioritizing insects recovered directly from banana inflorescences. Field-collected insects were surface sterilized, subjected to bacterial isolation, and confirmed by PCR using RSSC-specific and phylotype-specific primers; representative thrips were then morphologically identified. Preliminary acquisition assay with infected flowers for 1 and 3 days was conducted using field collected *T. hawaiiensis* from a non-Bugtok infested farm in Laguna. Among the insects recovered, thrips and stingless bees (*Tetragonula* spp.) were the most prominent flower visitors, but only thrips yielded internal detection of *R. solanacearum* after surface sterilization and homogenization, supporting the presence of the pathogen within the insect body rather than simple external contamination. Morphological characters of the positive specimens were consistent with *T. hawaiiensis*, including a pale antennal segment III, bicolored body, paired pronotal posteroangular setae, discal setae on abdominal sternite VII, and a complete comb on abdominal tergite VIII. In preliminary acquisition tests, healthy *T. hawaiiensis* exposed to infected flowers acquired the pathogen at mean positivity values of 1.75 ± 1.50 after 1 day and 2.67 ± 4.72 after 3 days, whereas control thrips remained negative. These findings provide field-based evidence that *T. hawaiiensis* can acquire *R. solanacearum* from infected *Saba* flowers and should be considered in Bugtok epidemiology and integrated disease management in Luzon.

## I. Introduction

Bugtok is a bacterial disease of cooking bananas in the Philippines, found in cultivars such as *Saba* and *Cardaba*, and is associated with members of the *Ralstonia solanacearum* species complex that invade the male inflorescence, rachis, and fruits, causing vascular discoloration and internal browning (Paret et al., 2024; Santos et al., 2020; Cellier et al., 2022) . Recent molecular work has reaffirmed that Bugtok belongs within the broader Moko ecotype of banana-infecting *Ralstonia*, highlighting the need for vigilant surveillance and improved understanding of local transmission pathways in Philippine production systems. In Luzon, concern over Bugtok has intensified because of its recent confirmed occurrence in Luzon, including recent outbreaks that started in Palawan in 2019, that later spread in Cavite in 2023. This later spread in a nearby province Batangas later that 2023, and now has reported incidence in a nearby area in Laguna in 2026, threatening the Saba-producing areas in Luzon . These alarming spreads highlight the need for an urgent investigation of local transmission pathways (Dela Cueva et al., 2023).

*Thrips hawaiiensis*, a flower-inhabiting, polyphagous thrips species attracted to floral tissues and volatiles, are known to be abundant on developing banana buds and flowers where they feed and move actively among inflorescence structures (Li et al., 2018; Wang et al., 2020). Because Bugtok infections are closely linked to floral entry points, a flower-feeding insect that repeatedly contacts infected ooze, nectar, and wounded tissues can function as a facilitator of pathogen acquisition and dissemination.

Related work from Southeast Asia strengthens this epidemiological hypothesis. In Indonesia, several flower-visiting insects, including stingless bees, common fruit flies (*Drosophila* spp.), earwigs, ants, and other banana inflorescence visitors, have been shown to carry *Ralstonia* associated with banana blood disease, and insect-mediated movement has long been recognized as important in the spread of banana bacterial wilt through floral pathways (Pinaria et al., 2020; Tangapo et al., 2019). However, direct evidence for *R. solanacearum* within *T. hawaiiensis* from Philippine *Saba* farms has been lacking, creating a critical knowledge gap addressed by the present study.

## II. Methodology

### Field collection and pathogen detection

Destructive field sampling was conducted three times, mid-morning under sunny conditions, in a Bugtok-infested farm of banana cv. *Saba* in Indang, Cavite, Philippines, within a PAGASA Climate Type I zone. Sampling was carried out from May to December 2025, spanning the late dry season, the wet season transition, and the early months of the northeast monsoon period, which together characterize the prevailing field conditions during the study period. The condition of each sampled plant was described as either healthy or infected with Bugtok based on visible disease symptoms present in the peduncle or its fruits. Because flowering plants were not randomly distributed across the farm at the time of assessment, sampling was opportunistic and prioritized inflorescences available during each collection event. Flowers were detached from the bearing plant using an elongated scythe and placed immediately into sterilized sampling bags. Insects associated with the inflorescence were collected using a fine brush, targeting specimens found on the bracts and florets, as well as those that dropped into the sampling bag upon flower detachment. All collected insects were transported to the laboratory for further analysis

### *R. solanacearum* isolation

All field-collected insects were subjected to a sequential surface sterilization procedure prior to homogenization. Insects were immersed in 10% sodium hypochlorite for 30 seconds then in 70% ethanol for 30 seconds and rinsed three times in sterile distilled water. Following sterilization, insects were individually homogenized in 100ul sterile distilled water. Homogenates were plated in a semi-selective media, tetrazolium chloride agar at room temperature for 72 hours. Colonies exhibiting white, fluidal, irregular morphology with pink centers were selected and then subcultured in the same media to be used for DNA extraction and PCR.

### DNA extraction

Putative colonies consistent with the *Ralstonia solanacearum* species complex were subjected to genomic DNA extraction following the modified SDS-based protocol adapted from Chen and Kuo (1993). A loopful of bacterial cells from a 48-72 h single colony was suspended in a 400ul of SDS-based lysis buffer, added with 150ul of 5M NaCl. The solution was vortexed and centrifuged at 14,000 × g for 15 minutes at 4 °C. The supernatant was combined with 400 μL of chloroform–isoamyl alcohol (24:1), mixed by inversion, and centrifuged at 14,000 × g for 10 minutes. The upper aqueous phase was transferred to a clean tube and precipitated with 2 volumes of ice-cold absolute isopropanol and incubated at -20 °C for 1-4 hours. This was followed by centrifugation at 12,000 × g for 10 minutes. The pellet was washed twice with 500 μL of 70% ethanol, air-dried in a laminar flow hood, and resuspended in 50 μL of TE buffer (10 mM Tris-HCl, 1 mM EDTA, pH 8.0) containing RNase A (100 μg/mL). DNA was stored at −20 °C until use.

### PCR-based identification

Isolates were initially confirmed as RSSC by PCR using the species-specific primers 759/760 (Opina et al., 1997). PCR reaction was prepared in a total volume of 15ul, containing 1× PCR buffer, 2 mM MgCl_2_, 0.2 mM dNTP mix, 0.2 μM of each primer, 1U Taq DNA polymerase, 5% dimethyl sulfoxide (DMSO), 2.5% bovine serum albumin (BSA), and 50 ng of DNA template. Thermal cycling consisted of initial denaturation at 96 °C for 5 minutes, 35 cycles of 94 °C for 15 seconds, 59 °C for 30 seconds, and 72 °C for 30 seconds, followed by a final extension at 72 °C for 10 minutes.

Subsequently, phylotype assignment was performed using the multiplex PCR (Fergan & Prior, 2005) using the 759/760 species-specific primers with the Nmult phylotype-specific primers. The reaction composition and thermal cycling parameters were as described above, with the exception that all seven primers were included simultaneously at equimolar concentrations of 0.2 μM each.

### Morphological identification of positive insects

All collected insect specimens were sorted and grouped by insect orders. Specimens associated with positive bacterial detection were subjected to morphological identification. Thrips specimens were identified to species level using the photo-based dichotomous key of Cluever and Smith (2017), with diagnostic characters examined under a stereomicroscope. Key morphological features assessed included antennal segment coloration, body color patterns, pronotal setal arrangement, and abdominal structural characters.

### Acquisition assay

To determine whether healthy *T. hawaiiensis* could acquire *R. solanacearum* from direct exposure to infected floral tissues, a laboratory acquisition assay was conducted using naturally infected *Saba* flowers. Thrips were sourced from the banana orchard of Plant Pathology Laboratory, Institute of Plant Breeding, University of the Philippines Los Baños – a site with no known history of Bugtok infection. Bugtok infected flowers were obtained from the study site in Indang, Cavite while healthy flowers were used as controls. Detached banana florets were individually placed in sterile test tubes sealed with cotton plugs. Healthy adult thrips were introduced into each tube and allowed to feed on infected flowers for 1 or 3 days. Control thrips were exposed to healthy flowers for the same durations. Following each acquisition access period, thrips were surface sterilized and individually transferred into 0.6mL microcentrifuge tubes containing 100ul sterile distilled water. Each tube was vortexed and a loopful of the suspension was streaked onto TZCA plates to verify sterilization efficacy. The thrips were then homogenized using sterile micropestle and processed for bacterial isolation and PCR confirmation as previously described. Differences in *R. solanacearum* acquisition among treatments were evaluated using Fisher’s exact test at 95% level of significance.

Healthy *T. hawaiiensis* from the orchard of plant pathology laboratory of Institute of Plant Breeding of University of the Philippines Los Baños were used as the exposure population. These healthy thrips were introduced to infected banana flowers and allowed acquisition access periods of 1 day and 3 days. Control treatments consisted of healthy thrips maintained with healthy flowers. After each acquisition period, thrips were recovered and processed using the same isolation and PCR confirmation workflow used for field-collected insects.

### Insects collected from *Saba* flowers

Across the three destructive collections, 65 *Saba* flowers were sampled, of which 41 were classified as healthy (65%) and 24 as infected (35%). The most prominent insects associated with both healthy and infected flowers were thrips and stingless bees (*Tetragonula* spp.) , whereas ants and earwigs were less consistently represented (Table 1).

**Table 1.** Frequency of Insect Occurrence in Healthy and Infected Saba flowers.

| Insect group | Healthy flowers | Infected flowers | Remarks |
| --- | --- | --- | --- |
| Ants | 7 | 8 | Present in both flower conditions |
| Stingless bees<br>(*Tetragonula* spp.) | 5 | 0 | Common among healthy flowers in this sampling set |
| Thrips | 22 | 13 | Most prominent insect group |
| Earwigs | 2 | 4 | Low-frequency flower visitors |

This insect group is epidemiologically important because flower-visiting insects repeatedly contact the male bud and floral surfaces that serve as likely infection courts in banana bacterial wilt systems (Pinaria et al., 2020; Tangapo et al., 2019). The repeated occurrence of thrips on both healthy and infected flowers supports continued exposure opportunities within the farm environment.

### Internal detection of *R. solanacearum* in thrips

The most important finding was that only thrips, and not the other prominent flower visitor highlighted in the study, yielded evidence of *R. solanacearum* carriage within the body. Non-sterilized thrips were not positive, whereas the same thrips became positive after series surface sterilization indicating detection from internal tissues rather than contamination.

Mean number counts further showed that thrips were abundant in both healthy and infected inflorescences and that positive thrips were recovered from both categories. Percent positivity exceeded 10% in both healthy and infected flower groups (Table 2).

**Table 2:** Mean number of collected thrips and positive thrips with percent positivity.

| Flower status | Mean thrips<br>± SD | Mean positive<br>thrips ± SD | Percent<br>positive |
| --- | --- | --- | --- |
| Healthy | 52.33 ±<br>35.76 | 8.67 ± 13.22 | 16.56 |
| Infected | 45.60 ±<br>20.66 | 7.00 ± 8.22 | 15.35 |

PCR confirmation (Figure 1) showed amplification of RSSC diagnostic fragments consistent with *R. solanacearum* detection from thrips isolates, including the expected species-complex and phylotype-specific bands used in established diagnostic assays. This molecular evidence strengthens the inference that *T. hawaiiensis* can harbor the pathogen internally after field exposure.

**Fig 1.**
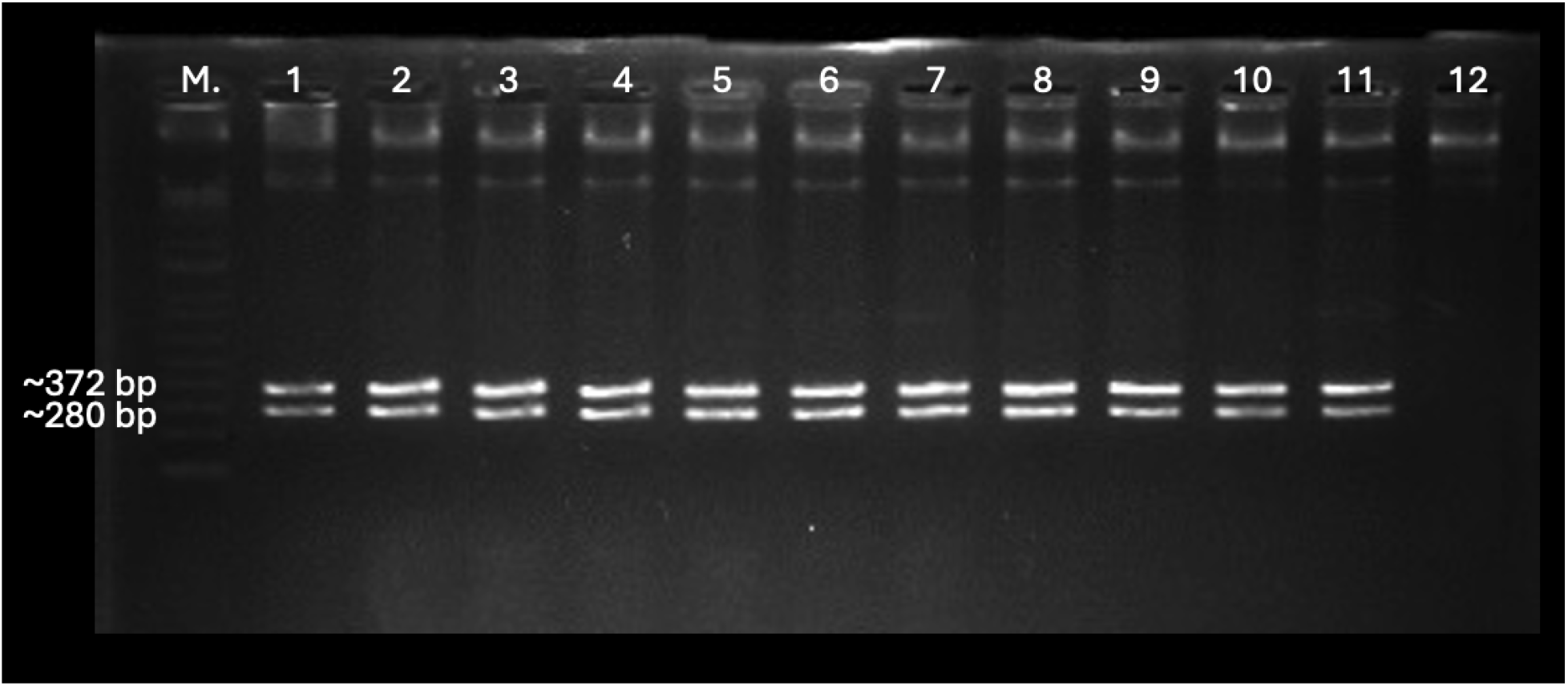
Agarose gel electrophoresis of multiplex PCR products from *R. solanacearum* isolated from surface-sterilized thrips. Species-specific primer pair 759/760 (Opina et al., 1997) yielded ∼282 bp, and Nmult primers (Fegan & Prior, 2005) produced ∼372 bp, indicating phylotype 2.

**Fig 2:**
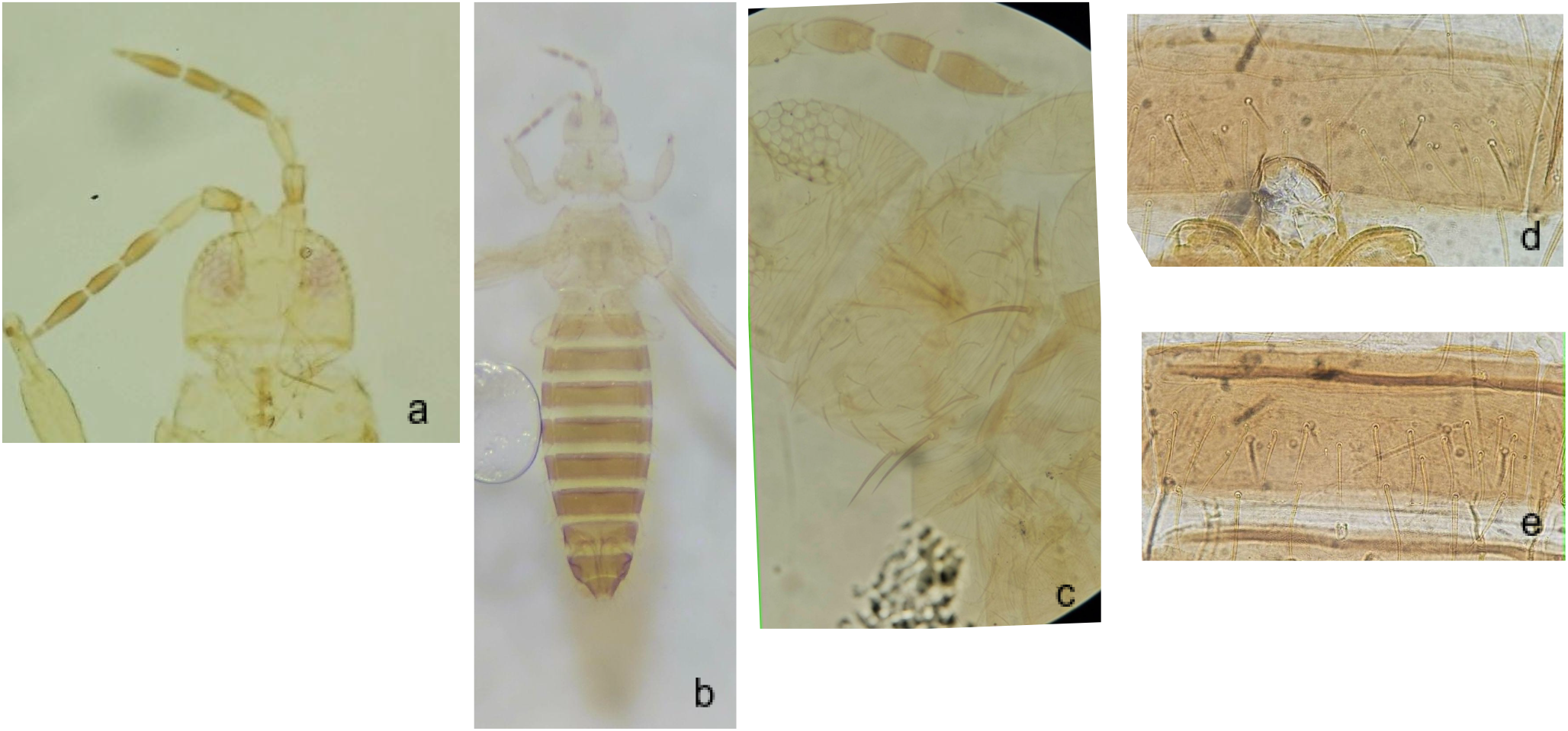
Diagnostic characters in positive thrips specimen from sampling in Indang, Cavite, a. Antennal segment III have yellow or light color than the rest of the segment, b. Bicolored body (Yellow to yellowish orange head and thorax and brown abdomen), c. Pronotum with 2 pairs of long posteroangular setae, d. Abdominal sternite VII with discal setae, e. Abdominal tergite VIII with marginal comb complete.

### Morphological identity of the positive insect

Morphological examination identified the positive thrips as *Thrips hawaiiensis* based on characters observed in the specimens. These features support the species assignment and are important for strong association of *T. hawaiiensis* with its known biology as a flower-dwelling thrips species (Li et al., 2018; Wang et al., 2020) . The key features emphasized and summarized below:

### Acquisition assay

In the acquisition assay, healthy *T. hawaiiensis* exposed to infected flowers showed measurable pathogen acquisition after both 1-day and 3-day exposure periods, whereas control thrips exposed to healthy flowers remained negative throughout the assay (Table 3). This pattern exhibits evidence beyond simple field association and demonstrates that healthy thrips can acquire the bacterium when allowed to expose and feed on infected floral tissues.

**Table 3:** Mean number and percent positivity of thrips following exposure to healthy and infected *Saba* banana flowers at different acquisition periods (1 and 3 days)

| Treatment | Acquisition period | Mean thrips ± SD | Mean positive thrips ± SD | Percent positive |
| --- | --- | --- | --- | --- |
| Infected flower | 1 day | 13.50 ± 6.19 | 1.75 ± 1.50 | 13.36% |
| Control healthy flower | 1 day | 10 | 0 | 0% |
| Infected flower | 3 days | 13.17 ± 7.63 | 2.67 ± 4.72 | 13.83% |
| Control healthy flower | 3 days | 12.33 ± 13.58 | 0.00 ± 0.00 | 0% |

Although the numerical increase between 1 day and 3 days was not interpreted as statistically significant, the higher mean positive count after 3 days suggests that extended contact with infected flowers may increase acquisition opportunities. At minimum, the complete absence of positives in the controls indicates that the assay signal is linked to infected-flower exposure rather than background contamination.

## III. Discussion

The present study provides a strong case that *Thrips hawaiiensis* is involved in Bugtok epidemiology in *Saba* banana in Indang, Cavite. The evidence rests on three associated observations: 1. Thrips were among the dominant insects associated with *Saba* flowers, 2. *R. solanacearum* was detected from within surface-sterilized and homogenized thrips, and 3. Healthy thrips acquired the pathogen after exposure to infected flowers under experimental conditions. Altogether, these findings support the statement that *T. hawaiiensis* is at least a biological facilitator of pathogen movement in the banana inflorescence environment and a possible candidate vector of Bugtok disease.

The internal detection result is especially important in insect-pathogen studies. *R. solanacearum* was only detected after sterilization and homogenization of the samples implies that the bacterium persists inside the insect body or alimentary tract and may be retained internally after feeding or close contact with infected flower tissues and not through accidental contamination from ooze or plant exudates (Pinaria et al., 2020; Tangapo et al., 2019). The acquisition assay strengthens this interpretation by showing that healthy *T. hawaiiensis* can become positive after only 1 to 3 days of exposure to infected flowers, while controls remain negative. This mirrors the biology reported for other banana bacterial wilt systems, where flower visitors contribute to pathogen spread by moving among male buds, floral bracts, nectar-rich tissues, and wounds that permit bacterial entry (Pinaria et al., 2020; Tangapo et al., 2019). Other studies also suggest that thrips can carry and acquire gut bacteria (Manday et al. 2025; Vries et al. 2001). In Indonesia, multiple banana flower visitors have been found to carry *Ralstonia* associated with blood disease, including stingless bees and other insects, demonstrating that floral visitation can be crucial to bacterial movement in banana agroecosystems (Montong & Salaki, 2019).

For the Philippines, the present work is significant because direct evidence of *R. solanacearum* in *T. hawaiiensis* from *Saba* flowers has been lacking even though Bugtok has long been recognized as a floral-entry disease. Recently, local studies on thrips infecting garlic proved the ability of these insects to carry bacteria (Manaday et al. 2025). Recent molecular studies continue to place Bugtok within the Moko ecotype of banana-infecting *Ralstonia* and emphasize the importance of accurate diagnostics and local epidemiological characterization for disease management (Paret et al., 2024; Santos et al., 2020; Cellier et al., 2022). By linking a common flower-inhabiting thrips with internal bacterial detection and experimental acquisition, this study fills part of that epidemiological gap. At the same time, the findings should be interpreted carefully. Detection and acquisition do not yet prove full vector competence in the strict sense, because successful transmission to healthy banana plants under controlled conditions was not demonstrated in the present dataset. Therefore, the conclusion is that *T. hawaiiensis* is a confirmed internal carrier of *R. solanacearum* in this system and a strong candidate vector of Bugtok disease, with transmission capability still requiring a dedicated inoculation study.

These results have immediate implications for Bugtok management in Luzon. If thrips contribute to pathogen movement at flowering, integrated pest management control should not focus only on sanitation but also on reducing floral exposure and insect access to infected floral buds. Practices such as prompt removal of diseased inflorescences, male bud management, field sanitation, and closer surveillance of flowering-stage insect activity may help reduce within-farm spread (Blomme et al., 2017) . This is particularly relevant in smallholder *Saba* systems where overlapping flowering cycles can maintain a continuous bridge for pathogen acquisition and movement.

In conclusion, the study provides the first field-based evidence from Indang, Cavite that *Thrips hawaiiensis* associated with *Saba* banana flowers can harbor *Ralstonia solanacearum* internally and can acquire the bacterium from infected flowers after short exposure periods. While further transmission assays, quantitative bacterial load measurements, and localization studies are still needed, the data strongly support inclusion of *T. hawaiiensis* in Bugtok epidemiology and justify its consideration in future integrated disease management programs for *Saba* banana in Luzon and elsewhere in the Philippines.

## IV. Acknowledgement

The team would like to express our sincere gratitude to the DA-BAR for funding the project and the local government unit of Indang, Cavite for the continuing support to advance the science of integrated pest management for Bugtok disease. We would like to acknowledge our support staff; Loi Pamulaklakin, Roland Gecalao, Aaron Villamor, Eddie Bueta, Aeron Pia. And our farmer beneficiary Marianito Blanco.

## References

1. Blomme, G., Dita, M., Jacobsen, K. S., Pérez Vicente, L., Molina, A., Ocimati, W., Poussier, S., & Prior, P. (2017). Bacterial diseases of banana and enset: Current state of knowledge and integrated approaches toward sustainable management. Frontiers in Plant Science, 8, 1290. 10.3389/fpls.2017.01290

2. Cellier, G., de Oliveira, J. C. F., Ceresini, P. C., Brioso, P. S. T., & Rossato, M. (2022). Comparative genomics and phylogenomics of the Ralstonia solanacearum Moko ecotype and its symptomatological variants. Genetics and Molecular Biology, 45(5), e20220132. 10.1590/1678-4685-GMB-2022-0132

3. Cluever, J. D., & Smith, H. A. (2017). A photo-based key of thrips (Thysanoptera) associated with horticultural crops in Florida. Florida Entomologist, 100(2), 454–467. 10.1653/024.100.0208

4. de Vries EJ, Breeuwer JA, Jacobs G, Mollema C. The association of Western flower thrips, Frankliniella occidentalis, with a near Erwinia species gut bacteria: transient or permanent? J Invertebr Pathol. 2001 Feb;77(2):120–8. doi: 10.1006/jipa.2001.5009. PMID: 11273692.

5. Dela Cueva, F. M., De Torres, R. L., Codrez, W. P., Mendoza, J. V. S., Tiongco, R. L., Manzanilla, C. M. C., Lucela, L. M., & Buendia, M. C. M. (2023). Managing the risk of spread of banana bugtok disease caused by Ralstonia solanacearum in the island of Luzon, Philippines. AGRIS/FAO. https://agris.fao.org/search/en/providers/122430/records/66fa7b6e1717de47eb54c7b7.

6. Fegan, M., & Prior, P. (2005). How complex is the “Ralstonia solanacearum species complex”? In C. Allen, P. Prior, & A. C. Hayward (Eds.), Bacterial wilt disease and the Ralstonia solanacearum species complex (pp. 449–461). APS Press.

7. Li, X., Fail, J., Shelton, A. M., & Feng, J. (2018). Effects of flower thrips (Thysanoptera: Thripidae) on nutritional quality of banana (Zingiberales: Musaceae) buds. PLoS ONE, 13(8), e0202199. 10.1371/journal.pone.0202199

8. Manaday, S. J. B., Reyes, C. P., Guerrero, M. S., Faroden, L. M., de Roxas, M. C. dL., & Caoili, B. L. (2025). Identification and characterization of bacteria isolated from garlic thrips (Thrips tabaci Lindeman) (Thysanoptera: Thripidae) in the Philippines. J. ISSAAS, 31(2), 41–53

9. Montong, V., & Salaki, C. (2020). Insects as carriers of Ralstonia solanacearum phylotype IV on Kepok banana flowers in South Minahasa and Minahasa districts. International Journal of ChemTech Research, 13(1), 199–205. 10.20902/IJCTR.2019.130124

10. Opina, N., Tavner, F., Hollway, G., Wang, J.-F., Li, T.-H., Maghirang, R., Fegan, M., Hayward, A. C., Krishnapillai, V., Hong, W.-F., Holloway, B. W., & Timmis, J. N. (1997). A novel method for development of species and strain-specific DNA probes and PCR primers for identifying Burkholderia solanacearum (now Ralstonia solanacearum). Asia-Pacific Journal of Molecular Biology and Biotechnology, 5(1), 19–30.

11. Paret, M. L., Alvarez, A. M., Allen, C., & Prior, P. (2024). Validation of PCR diagnostic assays for detection and identification of all Ralstonia solanacearum sequevars causing Moko disease in banana. Phytopathology. Advance online publication. 10.1094/PHYTO-06-24-0190-R

12. Pinaria, A. G., Assa, B. H., & Sembel, D. T. (2020). Insects as carriers of Ralstonia solanacearum phylotype IV on Kepok banana flowers in South Minahasa and Minahasa districts. International Journal of ChemTech Research, 13(1), 199–205.

13. Santos, A. S., de Oliveira, J. C. F., Ceresini, P. C., Brioso, P. S. T., & Rossato, M. (2020). Genomic sequencing of two isolates of Ralstonia solanacearum causing Sergipe facies and comparative analysis with Bugtok disease isolates. Genetics and Molecular Biology, 43(4), e20190362. 10.1590/1678-4685-GMB-2019-0362

14. Tangapo, A. M., Mantiri, F. R., Pelealu, J., & Sembel, D. T. (2019). Serangga pengunjung bunga pisang Kepok di Kabupaten Minahasa Selatan sebagai pembawa Ralstonia solanacearum filotipe IV (penyebab penyakit darah pisang). Eugenia, 25(3).15. Wang, Z.-H., Li, X.-W., Yang, X.-B., Zeng, Y., Huang, J.-R., Liu, W.-X., & Lu, Y.-Y. (2020). Behavioral responses of Thrips hawaiiensis (Thysanoptera: Thripidae) to volatile compounds identified from Gardenia jasminoides Ellis (Gentianales: Rubiaceae). Insects, 11(7), 420. 10.3390/insects11070420

